# Dynamic arousal signals construct memories of time and events

**DOI:** 10.1101/765214

**Authors:** David Clewett, Camille Gasser, Lila Davachi

## Abstract

Everyday life unfolds continuously, yet we tend to remember past experiences as discrete event sequences or episodes. Although this phenomenon has been well documented, the brain mechanisms that support the transformation of continuous experience into memorable episodes remain unknown. Here we show that a sudden change in context, or ‘event boundary’, elicits a burst of autonomic arousal, as indexed by pupil dilation. These boundaries during dynamic experience also led to the segmentation of adjacent episodes in later memory, evidenced by changes in memory for the timing, order, and perceptual details of recent event sequences. Critically, we find that distinct cognitive components of this pupil response were associated with both subjective (temporal distance judgements) and objective (temporal order discrimination) measures of episodic memory, suggesting that multiple arousal-mediated cognitive processes help construct meaningful mnemonic events. Together, these findings reveal that arousal processes may play a fundamental role in everyday memory organization.

## Introduction

As our lives unfold, we encounter a constant stream of sensory information. But our memories don’t strictly mirror the time and tide of experience. Instead, we remember the past as a series of discrete and meaningful episodes, or events. Furthermore, memory for the temporal duration of these episodes is subjective and often prone to distortion. For example, even if two experiences had occurred across the same objective amount of time, memory for their duration has been shown to be modulated by the content and structure of their constituent events [1-5]. While there has been intense interest in characterizing the factors that modulate this transformation from continuous experience into discrete episodic memories, little is known about the neural processes that support such memory-structuring, Thus, the aim of the present series of experiments was to address a critical question in learning and memory research: What brain mechanisms facilitate the creation of a new ‘episode’ in episodic memory?

Increasing research suggests that contextual stability over time plays a key role in integrating sequential information into memorable events. For instance, remaining in the same spatial context for an extended period of time, such as cooking breakfast in your kitchen, may help to organize a sequence of actions, such as cracking eggs and then frying them, into a unified event representation of ‘eating breakfast at home’ [6-8]. However, when the surrounding context changes, such as entering a new room or being interrupted by a phone call, people tend to perceive an ‘event boundary’ that defines the end of one event and the beginning of the next [8, 9]. Importantly, these event boundaries have a significant impact on how we remember these experiences later on, with boundaries leading to more separated memory representations [1, 4, 5, 10-19]. Thus, temporal stability and change in an unfolding context, including fluctuations in an individual’s surroundings or mental state, helps to form a mental timeline of discrete mnemonic events.

To date, the primary method of indexing the formation of discrete event memories has been to examine memory for the order and timing of sequential information, particularly when this information spans an intervening event boundary (for a review, see [11]). For instance, when individuals are presented with two items from a previously learned sequence, their temporal order memory is relatively impaired for items for which an intervening context shift occurred, such as a change in background color [13-20]. Likewise, individuals often remember items as having occurred farther apart in time if they had spanned an event boundary compared to items that had been encountered in the same context. These temporal memory distortions are subjective, given that the true temporal distance between these item pairs is actually equivalent [1, 4, 19]. Event boundaries have also been shown to influence non-temporal aspects of episodic memory, such as enhancing associative memory for an item and its local source information (e.g., background color; [17, 21, 22]). Thus, the existing literature presents a complex story, whereby temporal aspects of episodic memory integration are disrupted by changes in the surrounding context, while other ‘local’ aspects of memory (i.e., source binding) are enhanced.

We propose that one solution to this puzzle may involve fluctuations in physiological arousal—a notion inspired by evidence that spikes in arousal yield strikingly similar effects on episodic memory as do shifts in context. First, like event boundaries, emotional stimuli or acute stressors that induce arousal elicit exaggerated estimates of time duration [23, 24]. Second, viewing highly arousing videos prior to navigation or sequence learning has been shown to impair temporal order memory for neutral events [25-27]. Third, emotionally arousing stimuli also enhance local item-context source memory, at least when attention is directed towards relevant contextual information or when these details are intrinsic to the emotional stimulus [28-31]. This finding parallels the source memory enhancements observed for items that appear during an event boundary (e.g., [17]).

Importantly, fluctuations in arousal are induced by more than just emotion and stress. Many salient environmental changes, such as hearing an unexpected sound, can activate central arousal systems that regulate ongoing attention and memory processes across the brain [32-34]. Arousal signals also mediate many cognitive processes that are thought to be triggered by event boundaries, including increased cognitive control and prediction errors [8, 19, 34-39]. In light of this evidence, we hypothesize that arousal systems are ideally positioned to track the structure of unfolding experiences and to mediate the effects of context shifts on the discretization of events in memory.

In the current series of experiments, we tested this hypothesis by monitoring pupil size during a sequence learning task and examining if event boundaries trigger momentary bursts of arousal to promote event separation in memory. Participants encoded slideshows of 32 everyday objects displayed on a computer. To manipulate event structure during learning, we used a novel context manipulation of playing a simple auditory tone in participants’ left or right ear before each item. This tone remained the same for eight successive items, and then switched to the other ear to create an auditory ‘event boundary’, which served to parse the continuous 32-item sequence into four discrete sub-events with distinct auditory contexts. After each sequence, we then queried participants’ memory for the temporal order and temporal distance between the studied item pairs. Unbeknownst to participants, these pairs had always appeared the same objective distance apart during encoding. Critically, some item pairs were encountered in the same auditory event, whereas other item pairs spanned a boundary. Based on prior work, we predicted that these boundaries would lead to (1) relatively larger retrospective estimates of temporal distance between item pairs and (2) impaired temporal order memory. We also tested if boundaries enhanced participants’ auditory source memory for items appearing immediately after a tone switch (henceforth referred to as ‘boundary items’) relative to non-boundary items. Experiment 1 was performed to validate this paradigm.

In Experiments 2 and 3, we also measured pupil diameter continuously throughout our sequence task to index changes in autonomic arousal across time [40, 41]. Our primary goals were to determine whether event boundaries trigger robust spikes in arousal (measured as changes in pupil dilation), and whether these arousal changes were related to later memory separation effects, as evinced by changes in both objective and subjective aspects of episodic memory. For this we performed a temporal principal component analysis (PCA) on tone-evoked pupil dilations, an approach that has previously revealed distinct temporal features of the pupil response that may relate to different aspects of cognition [42-48]. In addition to targeting phasic arousal responses, this approach further enabled us to infer which mental processes (e.g., cognitive control, anticipation, etc.) that are modulated by boundaries also contribute to the separation of events in memory. Finally, we performed an exploratory trial-level analysis where we examined if moment-to-moment variability in pupil diameter over more prolonged periods of time also influences the temporal structure of memory. We reasoned that, insofar as physiological arousal itself may be a strong feature of an internal contextual state, greater temporal stability in arousal levels during encoding would promote temporal memory integration.

## Behavioral Results

### Experiment 1

To examine how context shifts influenced behavior during encoding, we compared response times to boundary items and non-boundary items for an indoor/outdoor judgement that participants made for each item (Figure 1). As expected, participants were slower to judge objects appearing immediately after a tone switch compared to items appearing after a repeated, or same-context tone, t(33) = 4.06, p < .001 [CI: 36.37, 109.61] (Figure 3).

**Figure 1.**
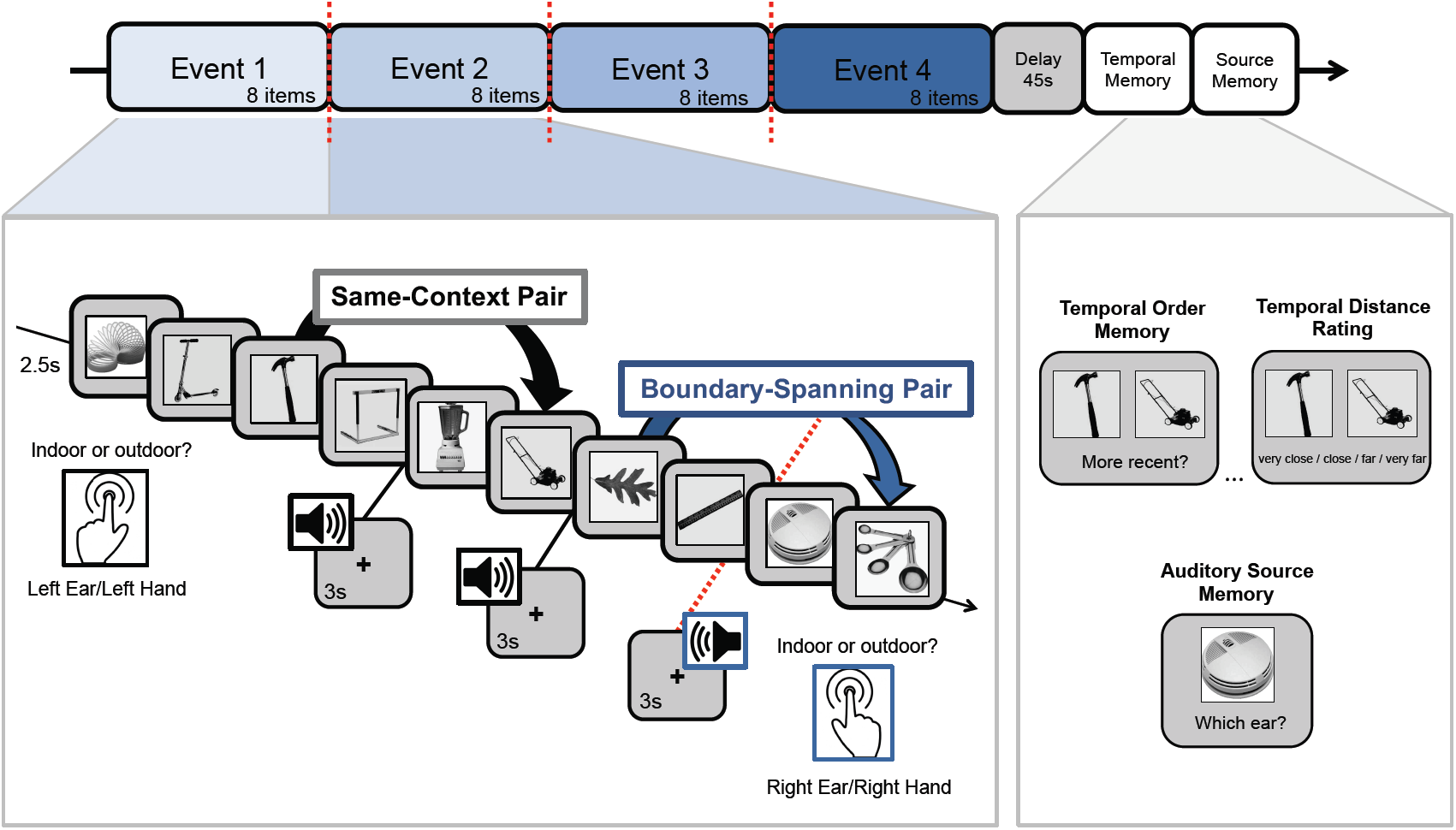
Auditory event boundary paradigm. Participants studied lists of 32 everyday objects and had to indicate whether each item would more likely be encountered in an indoor or outdoor setting. The surrounding context was manipulated by playing a simple tone in either participants’ left or right ear prior to viewing each image, which indicated to participants which hand they should use to make their subsequent indoor/outdoor judgement. After 8 successive items, the tone switched to the other ear and changed in pitch. These tone switches served as event boundaries, which parsed each continuous 32-item sequence into four discrete auditory events. After a short distractor task, participants performed three different memory tests in two separate blocks of trials: the first block included two different temporal memory tests and the second block included a source memory test. In the temporal memory block, participants first had to indicate which of the two presented items had appeared more recently in the prior sequence. Second, they had to rate the temporal distance between these items, ranging from ‘very close’ to ‘very far’ apart in the sequence. After being tested on item pairs, participants performed an auditory source memory test for all of the remaining items that were not shown during the temporal memory tests. In this source memory test, participants were shown individual items and had to indicate whether each item had been paired with a tone in their left ear or their right ear.

**Figure 2.**
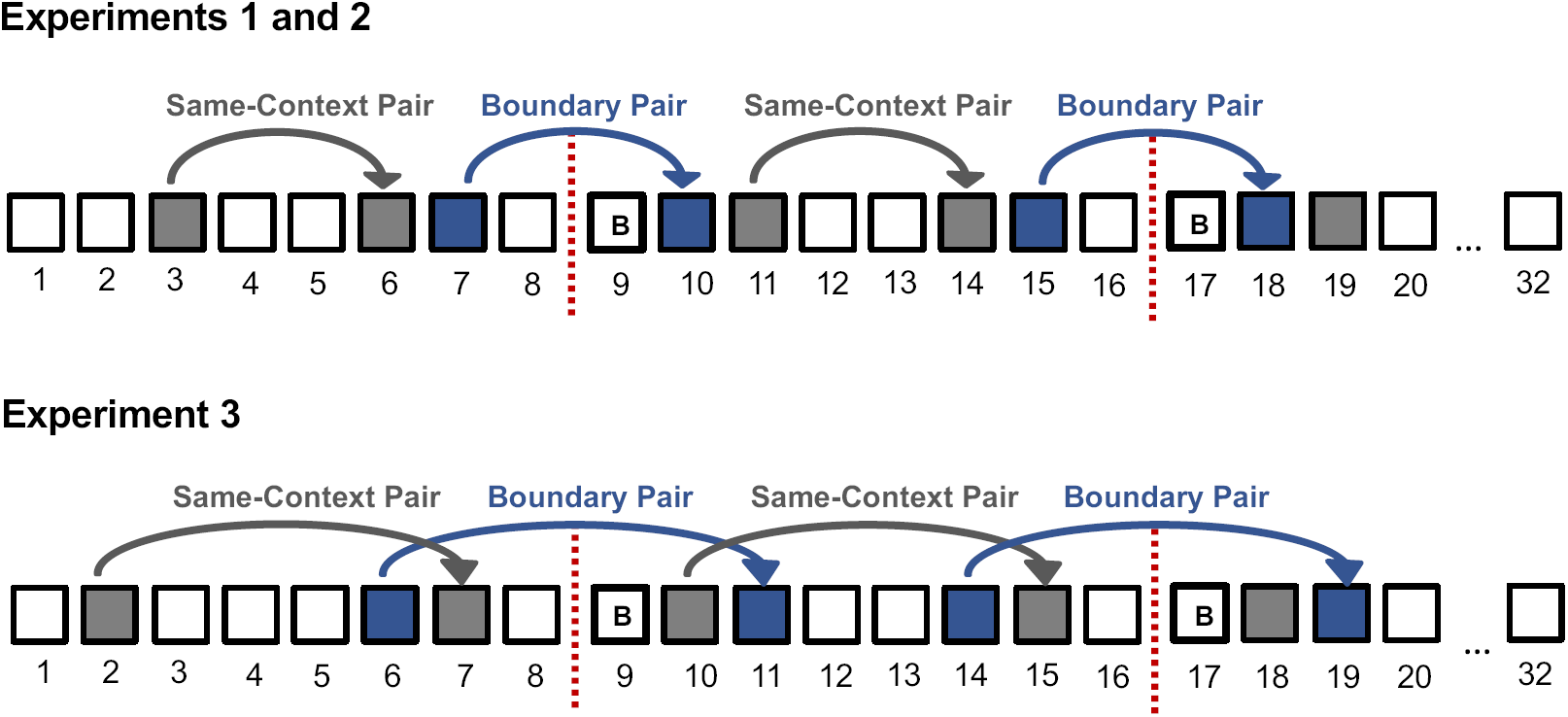
Schematic depicting the item positions for which different aspects of memory were tested after each 32-item slideshow. Colored boxes and arrows represent the positions of item pairs that were later tested in the temporal order memory and temporal distance memory tests. In Experiments 1 and 2, all of the tested item pairs had been presented with two intervening items during encoding. In Experiment 3, all of the tested item pairs had been presented with four intervening items during encoding. Dashed lines represent the location of event boundaries, or tone switches, which always occurred after 8 successive items. White boxes represent the positions of the individual items that were later tested for source memory (i.e., the ear that the accompanying tone played in). Boxes with a “B” signify boundary items, or the object images that appeared just after an event boundary. The remaining white boxes represent same-context items, except for the first and last items in each list, which were analyzed separately.

**Figure 3.**
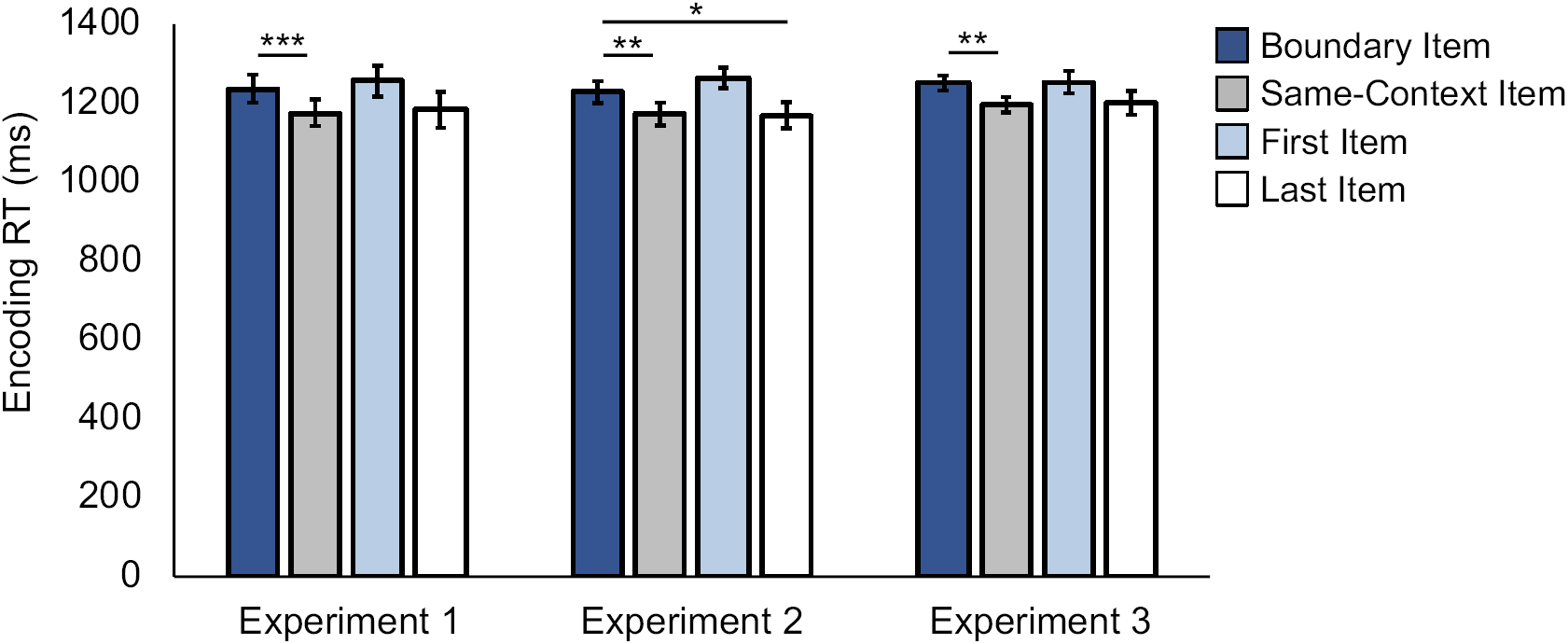
Mean response times (RTs) for indoor/outdoor item judgements during sequence encoding. Error bars represent standard error of the mean (SEM). ***p < .001; **p < .01; *p < .05.

Next, we examined how boundaries modulated the temporal structure of memory. We found that boundaries elicited a subjective time expansion effect in memory, such that boundary-spanning item pairs were later remembered as having appeared farther apart in time than same-context pairs, despite the objective distance being identical, t(33) = 2.44, p = .02 [CI: 0.015, 0.17] (Figure 4a). Temporal order memory (i.e., deciding which item in a pair appeared more recently) was significantly impaired for boundary-spanning pairs relative to same-context pairs, t(33) = −4.77, p < .001 [CI: −0.13, −0.052] (Figure 4b). Additionally, temporal order memory was significantly above chance for same-context pairs (p < .05) but not for boundary-spanning pairs (p > .05).

**Figure 4.**
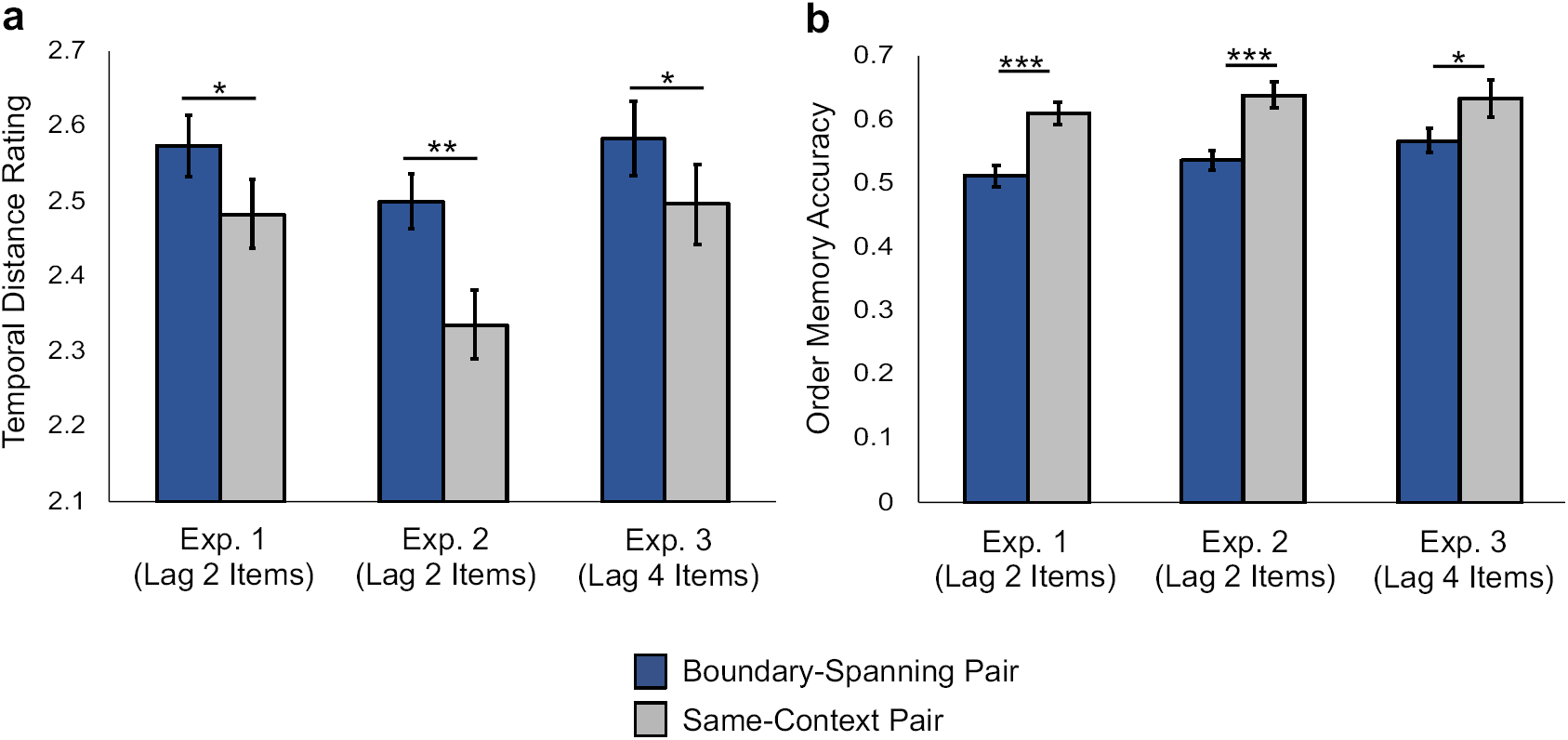
(a) Mean temporal distance ratings for item pairs from the object sequences. During this temporal memory test, participants rated how far apart the item pairs had appeared in the prior sequence, with choices ranging from ‘very close’ to ‘very far’. The ratings were then converted to a scale ranging from 1-4 and averaged together, such that higher values on the y-axis reflect more expanded retrospective estimates of temporal distance between item pairs. (b) Mean temporal order memory accuracy for item pairs from the object sequences. During this temporal order memory test, participants had to decide which of two items had appeared later (i.e., more recently) in the previous sequence of images. ‘Lag’ refers to the number of intervening items that had appeared between the to-be-tested item pairs at encoding. Error bars represent standard error of the mean (SEM). **p < .01; *p < .05.

Finally, source memory for the tone/ear paired with each item was significantly higher for items immediately following a boundary compared to those appearing in other event positions, F(3,25) = 8.68, p < .001, η^2^ = 0.51 (Figure 5). Specifically, source memory was enhanced for boundary items relative to items appearing in the same context, p < .001 [CI: 0.032, 0.12], as well as items from the very beginning, p = .025 [CI: 0.009, 0.15], and end of each list, p = .022 [CI: 0.023, 0.13].

**Figure 5.**
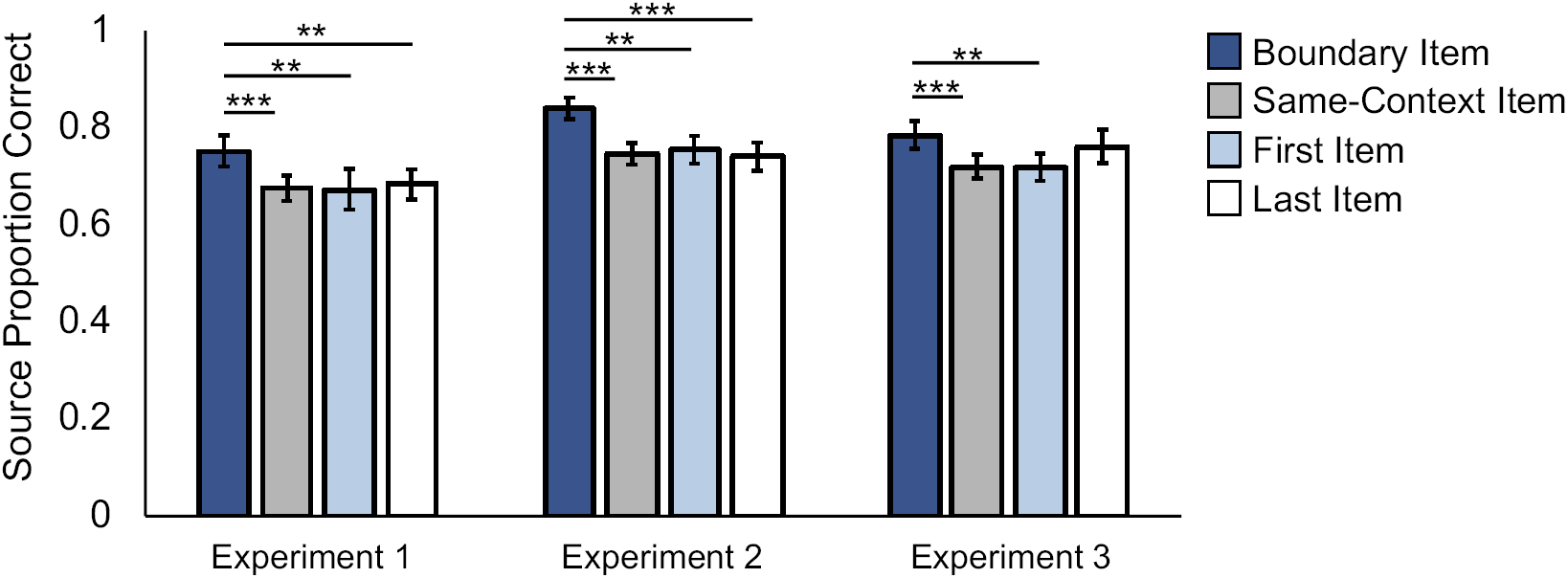
Mean source memory accuracy for individual items from the prior object sequence. During this test, participants had to indicate whether each individually-presented item had been paired with a tone played in his/her left ear or right ear during encoding. ‘Last Item’ refers to the last image presented in each 32-item list, whereas ‘First Item’ refers to the first image presented in each 32-item list. ‘Boundary Item’ refers to the first object appearing after a tone switch, or event boundary. Error bars represent standard error of the mean (SEM). ***p < .001; **p < .01; *p < .05.

### Experiment 2

The results from Experiment 1 validated our novel event boundary manipulation, showing that auditory context shifts elicit temporal and source memory effects consistent with the growing literature on how event memories emerge from continuous experience (for a review, see [11]). In Experiment 2, we combined this behavioral manipulation with eye-tracking to address our main hypothesis that fluctuations in physiological arousal during encoding, as indexed by pupil diameter, relate to the discretization of mnemonic events (described in the **Pupillometry Results** section below). All behavioral procedures were identical to those in Experiment 1.

Replicating the results from Experiment 1, participants were again slower to make the indoor/outdoor judgments at encoding for objects that followed a boundary compared to objects that appeared within the same auditory event, t(34) = 3.09, p = .004 [CI: 19.16, 92.55], and at the end of each list, t(34) = 2.55, p = .015 [CI: 12.00, 105.75] (Figure 3).

All of the memory results also replicated the findings from Experiment 1. Boundary-spanning item pairs were remembered as having appeared farther apart in time than same-context pairs, despite both item pairs appearing the same distance apart at encoding, t(34) = 3.18, p = .003 [CI: 0.06, 0.27] (Figure 4a). Temporal order memory was again worse for boundary-spanning item pairs compared to items from the same auditory context, t(34) = −6.45, p < .001 [CI: −0.14, −0.071] (Figure 4b). As in Experiment 1, source memory was significantly better for boundary items compared to items in other event positions, F(3,32) = 29.07, p < .001, η^2^ = .73 (Figure 5). Specifically, source memory was better for boundary items relative to items encountered within a stable auditory context, p < .001 [CI: 0.065, 0.12], as well as items at the very beginning, p = .004 [CI: 0.021, 0.15], and end of each list, p < .001 [CI: 0.046, 0.15

### Experiment 3

The behavioral results of Experiment 2 replicated the behavioral findings in Experiment 1, pointing to the robust effects of event boundaries on the organization of episodic memory. Prior event boundary experiments show that the lag between to-be-tested item pairs at encoding may influence subsequent temporal order memory performance [14, 49]. Thus, in the final experiment, we increased the objective distance between the to-be-tested item pairs from two to four intervening items (Figure 2) to examine if (1) the behavioral findings reported thus far are resistant to the actual distance between tested item pairs, and (2) potential relationships between arousal at boundaries and temporal memory are diminished (see **Pupillometry Results**).

Replicating the results from Experiments 1 and 2, participants were slower to make indoor/outdoor judgements for boundary items compared to items from a stable auditory context, t(29) = 3.15, p = .004 [C: 19.14, 90.17] (Figure 3). Participants again remembered boundary-spanning pairs as having appeared farther apart in time than same-context pairs, t(29) = 2.48, p = .019, [CI: 0.015, 0.16], despite, again, the actual distance being matched (Figure 4a). Temporal order memory was again worse for boundary-spanning pairs than for same-context pairs, t(29) = −2.72, p = .011, [CI: −0.12, −0.016] (Figure 4b). Also replicating the results from Experiments 1 and 2, source memory was significantly better for boundary items compared to other item types, F(3,27) = 8.56, p < .001, η^2^ = .49 (Figure 5), with memory being significantly higher for boundary items compared to items at other event positions (i.e., same-context items), p < .001 [CI: 0.027. 0.10] as well as for items appearing at the very beginning of each list, p = .03 [CI: 0.005, 0.13]. Source memory was not significantly better for boundary items compared to the last items in each list, p > .05 [CI: −0.044, 0.092].

## Pupillometry Results

Across all three experiments, we found reliable evidence that context shifts modulate both temporal and non-temporal features of episodic memories. To test our main hypotheses, we analyzed the pupil data from Experiments 2 and 3 to assess whether transient changes in arousal at event boundaries, as indexed by tone-triggered pupil dilation, promote the separation of adjacent experiences in memory.

### Pupil dynamics tracked event structure

Across both eye-tracking experiments (Experiments 2 and 3), fluctuations in pupil diameter appeared to be sensitive to event structure during encoding (Figure 6a). That is, although pupil size was dynamically modulated by the occurrence of *all* items and tones during sequence learning, transient spikes in pupil dilation were most robust at boundaries.

**Figure 6.**
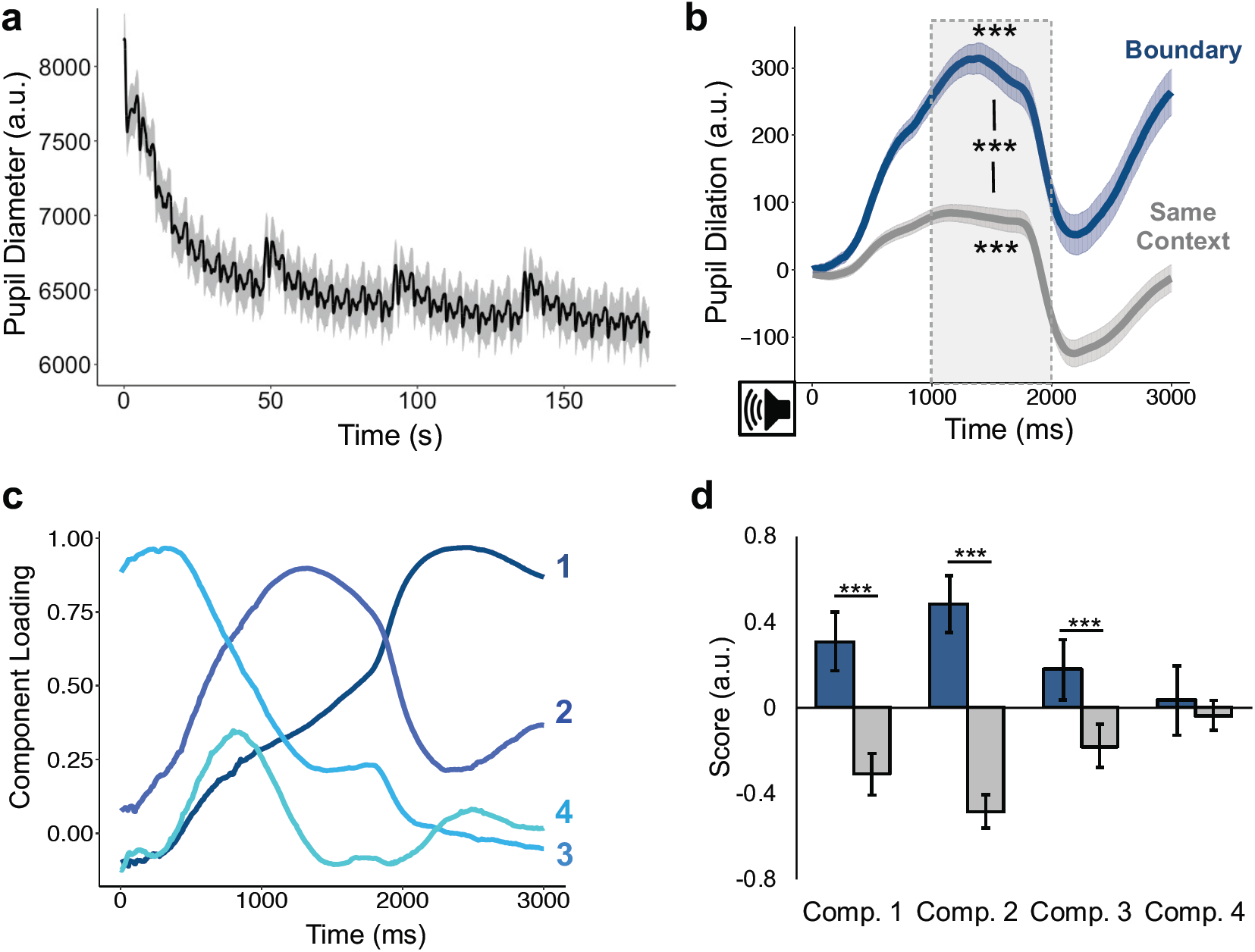
(a) Fluctuations in pupil diameter across the encoding sequence, averaged across all blocks, experiments, and participants. (b) The effects of boundary (tone switches; blue) versus same-context tones (repeated; gray) on pupil dilation. (c) Temporal features of tone-evoked pupil dilation identified by a temporal principal component analysis (PCA). The PCA characterized four significant features of pupil dilation that had distinct shapes over time. (d) The boundary tones (blue) versus same-context tones (gray) differentially modulated loading scores for the first three pupil components but not the fourth component (right panel). The gray boxes indicate the amount of variance in pupil dilation values that were accounted for by each component. All error bars represent standard error of the mean (SEM). ***p < .001; **p < .01; *p < .05.

To quantify the specific effect of boundaries on pupil size, we compared mean pupil dilation responses to the tone switches (boundary tones) versus pupil responses to repeated tones occurring within a stable auditory context (i.e., tones preceding items 2 through 8 in an event; Figure 6b). We first established that, across both eye-tracking studies, boundary tones, t(64) = 13.43, p < .001 [CI: 209.58, 282.84], and same-context tones, t(64) = 4.39, p < .001 [CI: 24.83, 66.34], elicited increased pupil dilation compared to baseline. Confirming our hypothesis, we also found that pupil dilation was significantly greater for boundary tones than for same-context tones, t(64) = 11.94, p < .001 [CI: 167.04, 234.20]. All of these results remained significant when analyzing data from Experiment 2 and Experiment 3 separately (all p’s < .05). These results demonstrate that auditory event boundaries were indeed salient and triggered a momentary increase in physiological arousal.

### Event boundaries modulate different temporal features of pupil dilation

Next, to further understand how boundaries modulate pupil dilation measures, as well as how this may impact ongoing cognitive processing, we performed a temporal principal component analysis (PCA) on the preprocessed pupil data. Prior research suggests that there are distinct temporal characteristics of pupil dilation that are regulated by different cognitive and neurophysiological processes, including cognitive control, motor responses, and salience detection [42-48]. Thus, this analytical approach enabled us to identify different components of pupil dilation, and to link ostensible cognitive components of pupil dilation to different episodic memory outcomes.

To identify pupil components, we averaged all of the pupil samples across the time-window of the tone-evoked pupil dilations (i.e., onset of tone plus three seconds; see Figure 6b) across participants and experiments. Importantly, the PCA was completely data-driven and agnostic to condition (i.e., boundary versus same-context tones). The PCA revealed four components that accounted for significant variance in tone-evoked pupil dilations (Figure 6c). The temporal features of these components, including their latencies-to-peak and variance accounted for, were as follows: (1) a late component (2,424 ms; 76.09% variance); (2) an intermediate component (1,316 ms; 16.35% variance); (3) a slowly-decreasing component (308 ms; 3.01% variance); and (4) an early-peaking component (800 ms; 2.66% variance). These pupil components were highly consistent with prior work applying PCA to pupil data, including a biphasic response that may signify separable contributions of the parasympathetic and sympathetic nervous systems to pupil size [45-48].

We next asked whether the degree to which individual participants ‘loaded’ onto these pupil components was modulated by event boundaries. Here, ‘loading’ refers to a measure of how much participants exhibited these distinct temporal patterns of pupil dilation in response to tones. Because we knew which data-points belonged to each condition, we were then able to compare loading differences between boundary and same-context tones. As shown in Figure 6d, boundaries significantly modulated loading values for the late component, t(34) = 4.81, p < .001 [CI: 0.31, 0.75], intermediate component, t(34) = 6.00, p < .001 [CI: 0.67, 1.35], and slowly-decreasing component, t(34) = 3.77, p = .001 [CI: 0.20, 0.68] across the two experiments. By contrast, there was no significant effect of boundaries on the early-peaking component loadings, t(34) = 0.29, p = 0.77 [CI: −0.43, 0.57]. The same results were also obtained when we examined the loadings from Experiments 2 and 3, separately (components 1, 2 and 4: p’s < .05; component 3: p’s > .05). Thus, boundaries significantly enhanced some but not all components of pupil dilation, suggesting that context shifts may engage specific mental processes and autonomic pathways.

### Spikes in arousal serve as a mechanism of memory separation

To test our key hypothesis that spikes in arousal contribute to event memory separation, we performed three across-subjects backward elimination multiple linear regression analyses modeling the relationships between the four pupil components and the three episodic memory outcomes: temporal distance ratings, temporal order memory and source memory.

Across both eye-tracking experiments, there was a significant positive association between pupil dilation at boundaries and temporal distance memory. Specifically, as shown in Figure 7a, individuals who exhibited a time expansion effect in memory for boundary-spanning pairs also showed more evidence of the early-peaking pupil component during boundaries versus non-boundaries (β = 0.36, p = .003). The relationship between the early-peaking pupil component and temporal distance memory was also significant in Experiment 2 (β = 0.35; p = .042) and Experiment 3 (β = 0.42; p = .020) when analyzed separately, suggesting that this relationship does not depend on the objective temporal distance between to-be-encoded item pairs at encoding.

**Figure 7.**
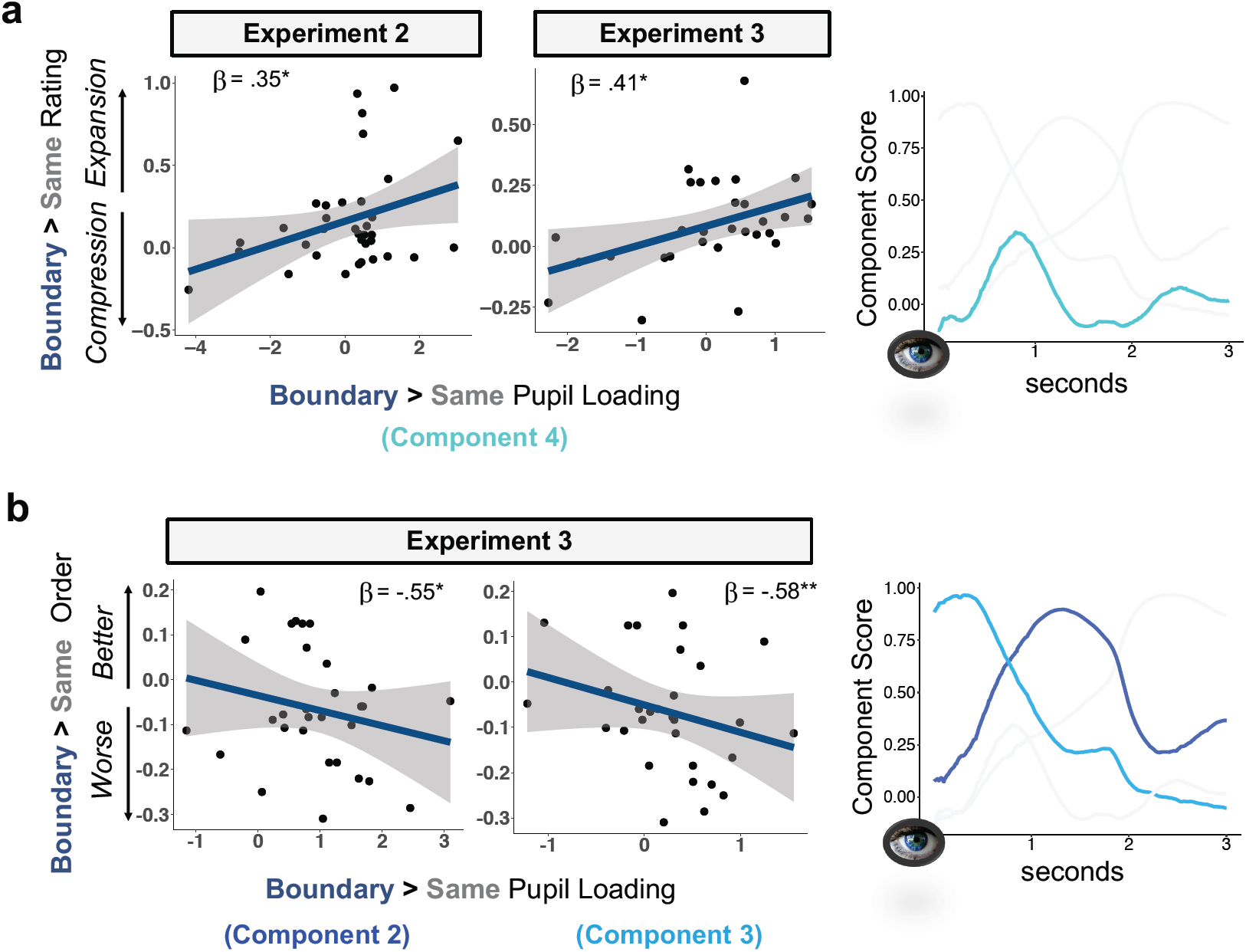
Relationships between different aspects of boundary-triggered arousal and temporal memory. (a) Association between boundary-modulated loadings on the early-peaking pupil component (#4; turquoise) and boundary-related effects on retrospective estimates of temporal distance between item pairs. Higher values on the y-axis reflect more expanded estimates of temporal distance for boundary-spanning pairs compared to same-context pairs. This pupilmemory relationship was significant in both of the eye-tracking studies (Experiments 2 and 3). (b) Association between boundary-modulated pupil loadings on an intermediate pupil component (#2; royal blue) and a slowly-decreasing pupil component (#3; light blue), and boundary-related effects on temporal order memory between item pairs in Experiment 3. Lower values on the y-axis reflect worse temporal order memory for boundary-spanning item pairs compared to same-context pairs. The bottom left plot represents the correlation between boundary-related temporal order memory and boundary-related loadings on the intermediate component. The bottom right plot represents the correlation between boundary-related temporal order memory impairments and boundary-related loadings on the slowly-decreasing pupil component. Gray shading in plots represents 95% confidence interval. **p < .01; *p < .05.

Next, we examined the relationship between the pupil dilation components and temporal order memory performance. We did not observe any significant pupil-memory associations when the data were collapsed across the two eye-tracking studies or in Experiment 2 alone. However, in Experiment 3, where the objective distance between the test pairs was the largest (i.e., four intervening items in Experiment 3 versus two in Experiment 2), we identified two significant relationships. As shown in Figure 7b, individuals who exhibited worse temporal order memory for boundary-spanning item pairs compared to same-context pairs also showed more evidence of the intermediate component (β = −0.55, p = .011) and the slowly-decreasing component (β = −0.58, p = .007) at boundaries relative to non-boundaries.

Additionally, there were no significant associations between any of the pupil components and enhanced boundary-item source memory (all p’s > .05).

Taken together, these results suggest that potentially unique cognitive and arousal processes, as indexed by distinct temporal characteristics of pupil dilation, modulate subjective and objective temporal aspects of episodic memory. These different pupil-memory relationships support our key hypothesis that brief increases in arousal at context shifts lead to more separated, or segmented, memories of temporally-adjacent experiences. Given the distinct relationships between our temporal memory outcomes and different cognitive and autonomic components of pupil dilation (e.g., [46]), our findings also suggest that multiple arousal-mediated mental processes may help construct event memories from continuous experience (discussed below). Interestingly, we did not observe any relationships between the pupil components and enhanced source memory at boundaries, suggesting that boundary-elicited arousal responses may be more important for modulating the temporal organization of events in memory.

### Temporal stability of arousal as a mechanism of memory integration

In an exploratory pupil analysis, we also tested if fluctuations in arousal over more prolonged periods of time support temporal memory integration. Behavioral evidence suggests that dynamic fluctuations in external contexts (e.g., space) over time modulate cognitive processes that link together sequences of information [11]. Here we reasoned that if arousal itself is considered an internal contextual ‘state’, its stability across time may be related to the integration of sequential representations in memory. Consistent with this idea, we found that lower trial-by-trial variability in pupil diameter between the to-be-tested item pairs at encoding (indexed by standard deviation in pupil size) was related to more compressed retrospective estimates of temporal distance and better temporal order memory (**Supplementary Materials**).

## Discussion

Addressing how the brain transforms continuous experience into memorable episodes is foundational to our understanding of learning and memory. Here, we provide evidence that (1) dynamic fluctuations in autonomic arousal across time are sensitive to the structure of unfolding experiences, and that (2) these arousal changes are in turn related to how those experiences become represented in memory. Specifically, we show that salient context shifts during sequence learning elicit increased pupil dilation. Furthermore, using a PCA approach, we found that dissociable temporal components of this arousal response – including those previously linked to increased cognitive control and response anticipation – were associated with temporal memory outcomes consistent with the discretization of events in long-term memory. Our findings underscore that arousal mechanisms during ongoing experience play a fundamental role in allocating memory representations to distinct events, with spikes in arousal at event boundaries contributing to the initiation and formation of temporally-distinct episodic memories.

A growing body of work suggests that temporal context stability (e.g., remaining in the same room for an extended period of time) may provide scaffolding for linking sequential experiences together in memory (for a review, see [11]). Conversely, a context shift may drive memories of adjacent experiences farther apart. Our findings lend support to this idea by showing that auditory event boundaries: 1) elicited greater subjective separation in memory between temporally-adjacent events; 2) impaired temporal order memory; and 3) enhanced source memory for local contextual features. These data expand upon earlier work using visual category or perceptual shifts as event boundaries by demonstrating that stability and change in an auditory context elicit similar memory integration and separation effects [1, 4, 5, 10-19]. Thus, many types of context shifts can drive event separation in memory, implicating a shared underlying process that is not limited to one sensory modality. Interestingly, we also demonstrate that the stability of arousal states over time helps integrate unfolding experiences into memorable events (see **Supplementary Materials** for results and discussion). This finding suggests that arousal is a strong feature of an internal contextual state that impacts memory organization in a similar manner to contextual information from the external world.

Using PCA to analyze our pupil data, we found that momentary increases in different aspects of pupil dilation at boundaries were associated with subjective and objective markers of event memory separation. The PCA first revealed prototypical temporal features of pupil dilation, which have been previously associated with different autonomic pathways that regulate pupil size [45, 46]. Furthermore, these different temporal characteristics of pupil dilation have also been linked to potentially dissociable aspects of cognition, such as sustained cognitive processing, mental resource allocation, and salience detection [42-48], suggesting that different mental processes are engaged by boundaries and promote later memory segmentation.

Across two eye-tracking experiments, we found that the early-peaking component of pupil dilation, which reflects parasympathetic regulation of pupil size under conditions that require increased mental effort and sustained cognitive processing [45, 46], was related to greater subjective time dilation effects in memory. Notably, this was the only component that was not significantly modulated by boundaries. This finding is intriguing, because it suggests that this pupil response may reflect a more generalized form of sustained cognitive processing that occurs as an experience unfolds. But *when* individuals show more of this arousal response to a boundary, they also seem to remember recent events as having occurred farther apart in time. One possible interpretation of this effect is that people are constantly maintaining an active representation of the world, or ‘event model’, which requires sustained cognitive processing [8]. At an event boundary, even greater cognitive control may be necessary to update this event model with new contextual information. This cognitive-updating process, in turn, may shift our memory systems to prioritize processing of a new context, creating greater separation in memory space.

Second, we linked boundary-related impairments in temporal order memory to both a slowly-decreasing and an intermediate pupil component, neither of which was associated with temporal distance ratings. The slowly-decreasing pupil dilation component (#3 in the current study) may reflect anticipatory arousal, given evidence that it is triggered in preparation of impending motor responses [43]. Moreover, this pupil component was also enhanced by event boundaries. In our experiments, all events were the same length, and there was an equal amount of time between each item in the sequences. As such, participants could likely predict when an impending tone switch would occur, as signified by an anticipatory pupil response. From this perspective, predictable changes in the environment could be used to proactively allocate ongoing experiences into contextually-appropriate memory representations. Indeed, individuals can still perceive discrete events even when the transition between them is predictable [50]. To further explore if anticipating a boundary during ongoing experience can guide event segmentation in memory, future studies could test whether this pupil-memory association disappears when context shifts are more or less predictable.

As discussed in the previous paragraph, worse temporal order memory across boundaries was also related to participants loading more on an intermediate pupil component at boundaries relative to non-boundaries. This pupil component peaked within a window typically attributed to mental resource allocation [41, 45], and has been shown to be regulated by sympathetic nervous system activation [46]. In prior work, the presence of this component has been related to reduced interference from previously encoded memories on the learning of new, conceptually-related information [43]. Thus, one interpretation for our finding is that reduced mnemonic integration across boundaries may stem from interference resolution between past and present context representations, especially if they share overlapping features. People also exhibit more evidence of this intermediate component when they hear salient oddball, or deviant, sounds compared to mundane sounds [48]. Emotionally arousing sounds also increase loading on this intermediate pupil component [47]. Together these findings raise the possibility that the salience, distinctiveness, and/or emotionality of a context shift may modulate the degree of mnemonic event separation for temporal-extended experiences. Importantly, our interpretations concerning the functional significance of these pupil components are drawn from reverse inference. Future work should thereby manipulate different cognitive processes at boundaries more directly (e.g., need for cognitive control), and examine how these changes relate to temporal components of pupil dilation and temporal memory outcomes.

Research in both animals and humans suggests that pupil dilation is a reliable biomarker of locus coeruleus-norepinephrine (LC-NE) system activity [51-54], which is known to modulate attention and the encoding of salient events [33, 34, 55]. LC neurons regulate pupil size both directly and indirectly through sympathetic and parasympathetic nervous system pathways, respectively [56, 57]. Thus, the functional neuroanatomy of the noradrenergic system makes it well positioned to support the observed boundary-related effects on temporal memory. Interestingly, core models of noradrenergic function propose that NE release helps to orchestrate a “network reset” that reorients attention and functional brain networks to process salient environmental changes [32]. This model bears a striking resemblance to a theoretical ‘reset’ signal that is thought to trigger event segmentation processes at boundaries [8, 58]. Furthermore, it is well established that under arousal, NE modulates memory processes in the hippocampus [59, 60], a brain region that is integral to binding contextual and temporal information in memory [61-63]. Taken together, our results suggest that arousal-related LC activity not only signals event boundaries, but also modulates processes that shape the temporal structure of memory downstream.

Of course, another possibility is that our results signify the contributions of multiple neuromodulatory systems to episodic memory organization. For instance, acetylcholine (ACh) release also co-occurs with pupil dilation [54] and regulates pupil size exclusively via parasympathetic nervous system pathways [56]. Given that we found a strong association between a parasympathetic-related feature of pupil dilation (the early-peaking component) and temporal distance ratings, one possibility is that ACh may specifically modulate subjective representations of time. Like NE, ACh also influences hippocampal encoding processes, particularly those that support the separation of experience into distinct memories [64]. Future work could investigate these neuromodulatory mechanisms using a combination of fMRI, eye-tracking and pharmacology.

The present findings have important implications for improving learning in both educational settings and everyday life. For instance, the ability to perceive and segment events is associated with enhanced memory for those events, even up to one month later [65-67]. The boundary-related impairments in temporal order memory we report might thereby be beneficial for structuring event memories in ways that improve long-term recall. Our work highlights that arousal fluctuations play an important role in this organizational process, and suggests that manipulating the structure of learning with salient and arousing event boundaries – as well as their associated mental processes – may be an especially effective strategy for enhancing long-term memory in the classroom and beyond.

## Methods

### Behavioral Methods: All Three Experiments

#### Experiment 1

##### Participants

Thirty-four individuals (23 women; M_age_ = 23.26, SD_age_ = 4.52) were recruited from the New York University Psychology Subject Pool and nearby community to participate in this experiment. All participants provided written informed consent approved by the New York University Institutional Review Board and received monetary compensation for their participation. A power analysis was performed on data from a similar event boundary experiment to estimate the appropriate sample size [14]. With an alpha = .05 and power = .80, we needed 28 participants to obtain a large effect size (d = .80; Cohen’s criteria [68]) for the temporal order memory effect (G*Power 3.1).

Additional participants were recruited in case of poor memory performance, potential withdrawal from experiment, or an inability to perform the task. We also expected overall temporal memory performance to be worse in the current experiment compared to the results reported in Dubrow and Davachi (2013), given that the sequence lists had eight additional items and there was a shorter lag between to-be-tested item pairs (two vs. three items). All eligible individuals had normal or normal-to-corrected vision and hearing, and were not taking beta-blockers or psychoactive drugs. For the source memory analysis, six participants were excluded from data analysis due to a programming error.

##### Materials

The object stimuli consisted of 480 images of everyday objects on a white background. These images were selected from previous datasets [69, 70]. Each image was resized to be 300 ×300 pixels and rendered in grayscale. To control for non-cognitive-related effects on pupil size, the luminance of all object images and fixation screens was normalized using the SHINE toolbox in MATLAB [71]. To manipulate the surrounding context during visual sequence learning, six 1s pure tones with sine waveforms of different frequencies (500Hz, 600Hz, 700Hz, 800Hz, 900Hz, 1000Hz) were generated using Audacity (https://www.audacityteam.org/). These frequencies were chosen such that sounds were discriminable from one another and were arousing enough to maintain participants’ attention.

##### Procedure

In the current study, we performed three separate behavioral experiments in which we queried different aspects of episodic memory from a sequence learning task. Building on prior studies from our lab, we developed a novel paradigm in which event boundaries within an image sequence were defined as a switch from one stable auditory context – in which the same tone was played in the same ear – to another (Figure 1).

##### Sequence encoding

For each sequence, participants viewed a series of 32 grayscale, luminance-normed images of objects. Each image was presented in the center of a gray background for 2.5 seconds. During the inter-stimulus interval (ISI) between each object, a black fixation cross was displayed in the middle of the screen for 3 seconds. Half-way through each ISI, or 1.5 seconds post-image, a 1-s pure tone was played in either the participant’s left ear or right ear. The ear that the tone played in indicated to participants which hand they should use to make their indoor/outdoor judgement (e.g., left ear = left hand). Specifically, the participant had to indicate via button press whether the displayed object would more likely be found in an indoor or outdoor setting.

Importantly, the specific tone/ear pairing heard before each object remained the same for eight successive object images, which served to create a stable auditory context, or ‘event’. After the 8^th^ item in each event, the tone switched to the other ear and changed in pitch, creating an ‘event boundary’. This new tone/ear pairing then remained the same for the next eight items, and so on and so forth. There were three event boundaries per list, creating a total of four auditory sub-events per list. The tone frequencies were pseudorandomized across lists, such that no tones of a given frequency were presented in more than one of the four sub-events within a list (i.e., in a given list, tones that were 700Hz were not heard in more than one 8-item event). The ear that the tones first played in was also counterbalanced across lists. Each participant viewed a total of 15 lists/sequences. The first list served as a practice block, allowing participants to become accustomed to the encoding and memory tasks. The data from the remaining 14 lists were included in all subsequent analyses.

##### Delay distractor task

To create a 45-s study-test delay, and to reduce potential recency effects, participants performed an arrow detection task after each sequence. In this phase, a rapid stream of either left-facing (<) or right-facing (>) arrow symbols appeared in the middle of the screen for 0.5s each. These arrow screens were separated by 0.5-s ISI screens with a centrally-presented black fixation cross. Participants simply had to indicate which direction the arrow was pointing via button press as quickly as possible.

##### Temporal memory tests

Following the distractor task, we tested three aspects of episodic memory. These tests were also divided into two different blocks of trials. The first block included two temporal memory tests, and the second block included source memory judgements. In the temporal memory block, participants were shown pairs of items from the prior sequence. First, we queried temporal order memory by having participants indicate which of two probe items from the prior sequence had appeared more recently (Figure 1). After this choice, the same pair of items remained on-screen, and participants had to rate the temporal distance between the two items. For this temporal distance rating, participants could rate item pairs as having appeared ‘very close’, ‘close’, ‘far’ or ‘very far’ apart in the prior sequence (e.g., [1]). Crucially, each pair of items had always been presented with two intervening items during encoding, and were thereby always encountered the same objective distance apart. Thus, any differences in temporal distance ratings between the two pair types were completely subjective. There was no time limit for each response. To test our hypothesis that event boundaries alter temporal memory organization, we considered two types of item pairs: (1) items that had appeared within the same auditory event (same-context pairs; 4 trials per list) and (2) items that had spanned an intervening tone switch (boundary-spanning pair; 3 trials per list).

##### Source memory test

In the second block of trials, we queried source memory for all of the items that had not appeared in the temporal memory test block (18 items total; Figure 2). Each item was displayed individually in the center of a gray screen. Participants had to indicate whether each object had been paired with a tone played in their left ear or in their right ear. There was no time limit for each choice, and the order of the items was re-randomized so that it did not match the presentation order from encoding. Examples of the item positions that were tested after each sequence are displayed in Figure 2 (white boxes).

##### Memory Analyses

Temporal order memory performance was calculated as the proportion of correct recency discriminations within each condition. To examine the effects of boundaries on order memory, these values were then submitted to a repeated-measures analysis of variance (rm-ANOVA; Context Condition: boundary-spanning pair, same-context pair). For temporal distance memory, the four possible ratings were converted to a scale ranging from 1 (‘very close’) to 4 (‘very far’) and then averaged together. The resulting distance values were submitted to a rm-ANOVA (Context Condition: boundary-spanning pair, same-context pair).

Source memory performance was calculated as the proportion of correctly remembered ear/sound side within each condition; specifically, whether participants successfully remembered whether an individual item had been paired with a tone played in the left ear or right ear. The source memory data was also analyzed using a rm-ANOVA, except this time the first item and the last item from each list were separated from the same-context condition to mitigate any potential recency or primacy effects on source memory (Context Condition: boundary item, same-context item, first item, last item). Planned follow-up two-tailed paired t-tests were then used to compare source memory accuracy for boundary items versus same-context items, as well for the first and last items in each list.

#### Experiment 2

##### Participants

Forty individuals were recruited from the New York University Psychology Subject Pool and nearby community to participate in this experiment. All participants provided written informed consent approved by the New York University Institutional Review Board and received monetary compensation for their participation. Inclusion criteria were the same as in Experiment 1. Five participants were excluded from data analysis: three participants withdrew mid-way through the experiment, one participant failed to follow task instructions, and the eye-tracker malfunctioned for one participant. Thus, data from thirty-five individuals were analyzed in this experiment (24 women; M_age_ = 22.57, SD_age_ = 4.24).

##### Procedure and memory analyses

The task for Experiment 2 used the same procedure as Experiment 1, with the addition of eye-tracking.

#### Experiment 3

##### Participants

Thirty-eight individuals were recruited from the New York University Psychology Subject Pool and nearby community to participate in this experiment. All participants provided written informed consent approved by the New York University Institutional Review Board and received monetary compensation for their participation. Inclusion criteria were the same as the first two experiments. A total of eight participants were excluded from data analysis: Four participants had poor eye-tracking quality (fewer than 50% valid samples) and four participants withdrew mid-way through the experiment. Thus, data from thirty individuals were analyzed in this experiment (21 women; M_age_ = 23.87, SD_age_ = 5.55).

##### Procedure and memory analyses

The task for Experiment 3 used the same procedure as Experiments 1 and 2, except for one modification. In this version, we presented four intervening items rather than two intervening items between the item pairs that were subsequently tested during the temporal memory tests (see Figure 2). All of the behavioral analyses were the same as in Experiments 1 and 2.

### Eye-Tracking Methods: Experiments 2 and 3

#### Eye-tracking

During sequence learning, participants were seated 55 cm from the computer and pupil size was measured continuously at 250 Hz using an infrared EyeLink 1000 eye-tracker system (SR Research, Ontario, Canada). By presenting tones during fixation screens appearing between the objects, we were able to acquire clean measures of pupil dilation that were unconfounded by stimulus brightness and visual complexity. The pupil data were preprocessed and analyzed using in-house code implemented in Matlab 9.4 (MathWorks, Natick, MA). Experiment blocks (sequences/lists) with less than 50% valid pupil data were removed from the final analysis. Eye-blinks and other artifacts, such as signal dropout, were removed using linear interpolation.

#### Pupil dilation analysis

Tone-evoked pupil dilations were compared for two events of interest: the boundary tone (i.e., tone switch after the 8^th^ item in an event) and the same-context, or repeated, tones (i.e., the seven subsequent tones that remained stable within an event). To measure pupil dilation responses, average pupil diameter was measured from 1-2s after cue onset when the tone-evoked pupil response was most apparent (Figure 6b). The average pupil size during this time-window was then baseline-normed by subtracting the average pupil size during the 500ms window prior to tone onset. To examine the influence of event boundaries on pupil dilation, we performed two-tailed paired t-tests comparing average pupil dilation responses between the boundary (i.e., tone switched) and same-context (i.e., tone repeated) trials.

#### Pupil dilation temporal principal component analysis (PCA)

To analyze how event boundaries altered different components of pupil dilation, we performed a temporal principle component analysis (PCA) by adapting methods described in Johansson et al. (2018). Experiments 2 and 3 were conducted in the same behavioral testing room and under the same dimly-lit lighting conditions. Thus, we combined the data from the two eye-tracking experiments to take advantage of additional statistical power to detect different pupil components. This also enabled us to draw comparisons between the same components across the two eye-tracking studies. Average baseline-normed pupil dilation was computed for each of the tone types (boundary and same-context trials). These values were then averaged across participants to reduce the amount of input variables (130 variables; 65 participants with two conditions each) and to control for noisy, spontaneous pupil changes on individual trials. All pupil samples across a 3-s window covering the average time-course of tone-evoked pupil dilation (750 pupil samples in total; see Figure 6b) served as dependent measures in the PCA.

A Varimax rotation was performed on the components output by the temporal PCA, which were defined based on an eigenvalue equal to or greater than the average value of the original variables. An unrestricted PCA using the covariance matrix with Kaiser normalization and Varimax rotation was used on all components to generate maximal component loadings on one component with minimal overlap with other components. The resulting loadings reflect the correlated, temporally dynamic patterns of pupil dilation elicited by the auditory tones.

Because the PCA was data-driven and agnostic to condition, we were able to then examine the relative contributions of loading patterns for each pupil component to same-context versus boundary tones. Two-tailed paired t-tests were performed on these loading values to determine how boundaries modulated different temporal characteristics of pupil dilation.

#### Relationship between pupil dilation components and memory organization

To test our key hypothesis that increased arousal at boundaries modulates episodic memory organization, we next performed three multiple linear regression analyses for each of the three memory outcomes (temporal distance, temporal order, and source memory) in Experiments 2 and 3. These were first performed by collapsing the data across both experiments. In order to see whether the effects replicated in the two eye-tracking experiments individually, we then analyzed the data from each experiment separately. Because we had no a priori predictions about which specific pupil components would relate to behavior, we performed two-tailed, backward elimination linear regressions.

## Supporting information

supplemental results/methods/figure/discussion

## Acknowledgements

This project was funded by federal NIH grant R01 MH074692 to L.D. and by fellowships on federal NIH grants T32 MH019524 and F32 MH114536 to D.C. The authors thank Alexander Ren for his assistance with data collection. We also thank Sarah Barber, Elizabeth Goldfarb, and Oded Bein for helpful comments on earlier versions of this manuscript.

