## supplemental results/methods/figure/discussion for "Dynamic arousal signals construct memories of time and events"

### Supplementary Materials

#### Supplementary Results

***Temporal stability of arousal serves as a mechanism of temporal memory integration.*** The pupil dilation results from our PCA analysis suggest that a spike in arousal at event boundaries can indeed promote event segmentation in later memory. In a more exploratory pupil analysis, we asked if prolonged fluctuations in arousal over time support temporal memory integration. Here we reasoned that if arousal itself provides information about the current context, it should also serve as a proxy for the stability of ongoing cognitive operations as experiences unfold. Through this lens, we predicted that greater stability in arousal levels over relatively long periods of time (e.g., 19+ seconds; as indexed by lower variability in pupil diameter changes) would promote stronger temporal memory integration.

To test this hypothesis, we used a separate analysis approach that enabled us to leverage the rich time-course of pupil data on the order of tens of seconds on a trial-by-trial basis. Specifically, we performed linear mixed effects regression analyses to examine the relationship between trial-by-trial variability in pupil diameter between the to-be-tested item pairs and the two subsequent temporal memory outcomes (see Supplementary Figure 1, top panel). A paired t-test revealed no main effect of event boundaries on pupil size variability ($p > .05$). This is perhaps unsurprising, given that the average tone-evoked pupil dilation lasted approximately three seconds (Figure 6b) – a window that covered only ~16% of the interval between to-be-tested item pairs in Experiment 2 and 10% of the time interval between item pairs in Experiment 3. Thus, we collapsed the data across conditions (boundary vs. same-context) to examine if arousal fluctuations – irrespective of context changes in the external world (i.e., repeated tones or tone switches) – were associated with our temporal memory outcomes.

We first queried the relationship between pupil size variability across time and an objective measure of temporal memory: temporal order memory. Across both eye-tracking experiments (Experiments 2 and 3), we found that lower pupil size variability during the time window between a given pair of items at encoding was associated with successful temporal order memory of those items ($z = -2.09$, $p = .037$; Supplementary Figure 1, bottom left). When the two eye-tracking experiments were analyzed separately, this relationship was significant in Experiment 2 ($z = -2.55$, $p = .011$), which had a shorter interval between to-be-tested item pairs. However, we did not observe this relationship in Experiment 3, which had a longer time interval between to-be-tested item pairs ($z = -0.23$, $p = .82$). Thus, at least at the shorter timescale examined in Experiment 2 (i.e., 19 seconds), it appears that arousal stability may support temporal order memory.

Next, we examined the relationship between pupil size variability across time and a subjective measure of temporal memory: temporal distance ratings. Collapsing across both Experiments 2 and 3, we found that lower pupil size variability between a given item pair at encoding was associated with more compressed retrospective estimates of temporal distance between that same item pair ($z = 2.05$, $p = .040$; Supplementary Figure 1, bottom right). This pattern emerged in both eye-tracking experiments individually, but was smaller and trended towards statistical significance in both Experiment 2 ($z = 1.48$, $p = .14$) and Experiment 3 ($z = 1.57$, $p = .12$) when analyzed separately.

Together, these results support the idea that greater temporal stability in arousal states across more prolonged periods of time provide a form of context that guides the binding of sequential representations in episodic memory. The results from the temporal order memory analysis suggest that arousal stability promotes the encoding and preservation of order information in memory. However, this temporal memory integration effect may only occur at relatively shorter timescales of encoding, as this relationship was not observed when item pairs were studied farther apart in the sequence (i.e., in Experiment 3).

The results from the temporal distance ratings analysis suggest that arousal stability may also facilitate the creation of tight-knit and temporally-compressed episodic memories. This pattern was consistent across both eye-tracking experiments, suggesting that the objective interval between two to-be-associated items might not moderate the strength of this subjective temporal memory effect.

#### Supplementary Discussion

Interestingly, our findings raise the possibility that internal contexts, or states, may guide memory integration in a manner similar to external changes in context, such as shifts in perceptual or spatial information. Using a trial-level analysis, we found that lower variability in pupil diameter across time, as measured by the standard deviation in pupil size during the interval between two items (on the order of 19 or 30 seconds, depending on which eye-tracking experiment) was associated with indices of mnemonic event formation; that is, evidence of improved temporal order memory and more compressed retrospective estimates of temporal distance between those items. Past work suggests that there are many organizational principles by which temporally-extended experiences cluster together in memory, including items' temporal context [1], semantic similarities [2, 3], and overlapping perceptual features [4-6]. However, this existing body of work mostly considers contextual information that is either theoretical in nature (e.g., mental context; [1, 7]) or comes from the external world (e.g., categories, colors, space etc.; [8]). Our data add to this growing body of research by demonstrating that fluctuations in physiological states may also provide a medium for integrating sequential information into memorable events.

#### Supplementary Methods

***Pupil diameter variability analysis.*** To measure the stability of arousal fluctuations across sequence learning, we measured the standard deviation in pupil diameter between the onset of the first item from a to-be-tested memory pair and the offset of the second item from that pair (see Supplementary Figure 1, top panel). This allowed us to leverage the rich, temporally-extended fluctuations in pupil samples between the time participants encountered these item pairs. Across the two eye-tracking studies, the temporal distance between the to-be-tested item pairs were as follows: Experiment 2 = 19 seconds (4,750 pupil samples) and Experiment 3 = 30 seconds (7,500 pupil samples).

To query the relationship between this trial-level pupil size variability measure and temporal memory, we performed separate hierarchical linear modeling (HLM) analyses using the glmer function in the lme4 library. Parameters were estimated with the maximum likelihood method in R (R Core Team, 2012). This approach is exceptionally sensitive to linear modulation of memory outcomes by trial-level variations in pupil diameter.

The standard deviation values of pupil sizes between the to-be-tested image pairs were mean-centered within each participant and modeled as level-1 predictors. Model fits included random intercepts and slopes for pupil variability (i.e., standard deviation) by participant, enabling us to capture individual differences in pupil-memory associations. Condition (boundary-spanning pair vs. same-context pair) was also entered as a fixed-effect predictor in each model.

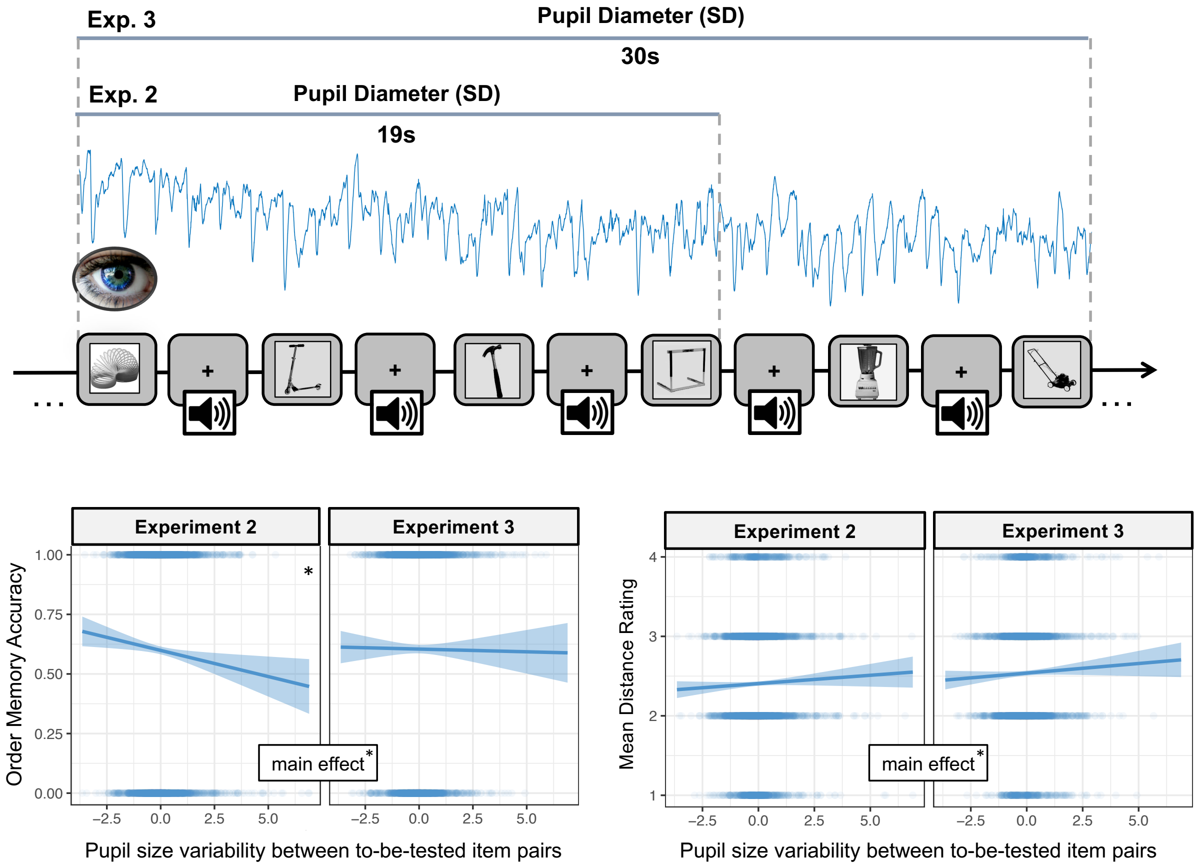


**Supplementary Figure 1.** The temporal stability of arousal states over relatively long periods of time facilitates temporal memory integration. (Top Panel) To assess variability in arousal fluctuations across each sequence, we measured the standard deviation (SD) of pupil diameter values across all of the pupil samples between to-be-tested item pairs. This time period was measured from the onset of the first image from a pair to the offset of the second image from that pair. In Experiment 2, the temporal distance between item pairs was 19 seconds, and in Experiment 3, the temporal distance between item pairs was 30 seconds. (Left Bottom Panel) Mixed effects linear modeling revealed a significant relationship between temporal recency discrimination and pupil size variability, such that participants were better at remembering the order of item pairs if there had been lower variability in pupil size between those item pairs at encoding. This linear pupil-memory effect was significant when data from both eye-tracking studies were collapsed. (Right Bottom Panel) Mixed linear regression analyses also revealed a significant linear relationship between temporal distance ratings and pupil size variability, such that participants were more likely to remember item pairs as having appeared closer together in time if there had been lower variability in pupil diameter between those item pairs at encoding. This pupil-memory effect was significant when data from both eye-tracking studies were collapsed. *$p < .05$.
